# Increased Mistranslation Protects *E. coli* from Protein Misfolding Stress due to Activation of a RpoS-dependent Heat Shock Response

**DOI:** 10.1101/698878

**Authors:** Christopher R. Evans, Yongqiang Fan, Jiqiang Ling

## Abstract

The misincorporation of an incorrect amino acid into a polypeptide during protein synthesis is considered a detrimental phenomenon. Mistranslated protein is often misfolded and degraded or non-functional and results in an increased cost to quality control machinery. Despite these costs, errors during protein synthesis are common in bacteria. Here we report that increased rates of mistranslation in *Escherichia coli* provide protection from protein misfolding stress by increasing the level of the heat shock sigma factor, RpoH. Surprisingly, this increase in RpoH due to mistranslation is dependent on the presence of the general stress response sigma factor, RpoS. This report provides evidence for a protective function of mistranslation and suggests a novel regulatory role of RpoS on the RpoH-activated heat shock.

## Introduction

Errors during protein synthesis can be the result of external stresses, such as aminoglycoside treatment [1], or failures in proofreading and protein quality control [2]. Each of these mechanisms result in the production of proteins with incorrect amino acids within the polypeptide. These mistranslated proteins often result in non-functional and misfolded protein products and an inherent cost to quality control mechanisms within the cell. Under severe mistranslation stress, this can ultimately result in growth inhibition or cell death [3, 4].

In order to handle the stress of misfolded protein accumulation, bacteria activate a highly conserved alternative sigma factor, sigma 32 (RpoH) [5, 6]. RpoH production and activation is complex and highly regulated due to the costly nature of the heat shock response [5-8]. RpoH activation results in the production of proteases and chaperones, including DnaK and GroEL, that collectively refold or degrade misfolded proteins [7]. Once the misfolded protein pool has been appropriately managed, these proteases degrade RpoH, resulting in a dynamic and transient regulation of the heat shock response [9-12].

Interestingly, natural isolates of *E. coli* have a wide range of ribosomal error-rates, indicating that, despite their cost, errors during translation may be beneficial under some conditions [13]. Recent studies have found examples of direct mechanistic benefits of mistranslation and broad physiological changes due to mistranslation that can be beneficial to cells [14-18]. In the case of mechanistic benefits, increased methionine misincorporation into proteins can increase oxidative stress tolerance by sequestering oxygen radicals away from functional proteins [19-21]. Also, increased mistranslation is sufficient to generate gain-of-function heterogeneous protein pools that can be resistant to antibiotic binding [15]. Physiological changes to cells as a result of mistranslation can protect cells by pre-activating stress responses. For example, increased mistranslation rates activate the general stress response sigma factor, RpoS, which protects the cells from future lethal oxidative stress [17, 22].

In this report, we show that RpoH is activated in *E. coli* cells with a moderately increased mistranslation rate. This response allows cells to survive lethal heat stress and manage misfolded protein stress better than cells with a lower mistranslation rate. Surprisingly, this protective heat shock response is dependent on the general stress response sigma factor, RpoS. This indicates a novel regulatory role for RpoS for mistranslation-induced heat shock response activation and a second benefit RpoS provides due to increased mistranslation rates.

## Materials and Methods

### Bacterial strains and growth conditions

*E. coli* cultures were grown in Luria broth (LB) and plated onto Luria agar plates. Cultures were grown at 30 °C shaking unless otherwise indicated. The following antibiotics were used for plasmid maintenance: 100 μg/mL ampicillin, 25 μg/mL chloramphenicol.

### Heat killing assay

*E. coli* cultures were grown overnight in LB from individual colonies then diluted 1:100 or 1:50 (Δ*rpoS* strains) in LB. The diluted cultures were grown for 3 hours at 30 °C to mid-logarithmic phase. To test the effect of canavanine on heat killing, MG1655 cultures were grown in LB supplemented with 3 mg/mL canavanine for one hour. To control for the decrease in growth due to canavanine treatment, separate MG1655 cultures were treated with 0.5 μg/mL chloramphenicol. Then, 500 μL of the cultures were centrifuged at 4000 rpm for 8 minutes and washed with phosphate buffer. After a second wash, the cultures were resuspended in 500 μL of phosphate buffer. The absorbance (A600) of the cultures in phosphate buffer was determined via spectroscopy in a microplate reader (Synergy HT, BioTek) and normalized to A600 = 0.5. After normalization, the cultures were further diluted to A600 = 0.1 and placed in a 96-well plate (Corning). Samples were subjected to 50 °C heat shock in a water bath. A 20 μL aliquot of each sample was taken every 30 minutes, and 5μL of serial dilutions was spotted onto an LB agar plate, and the survival were determined by the colony forming units (cfu).

### Fluorescent aggregation assay and image analysis

*E. coli* strains were grown overnight in LB with 100 μg/mL chloramphenicol. Then, they were diluted 1:100 in LB with 100 μM isopropyl β-D-1-thiogalactopyranoside (IPTG) to induce *sfGFP-ClpB* expression and grown for 3 hours at 30 °C. To induce aggregation, mid-logarithmic phase cultures were treated with 100 μg/mL streptomycin or incubated at 42 °C for one hour. After treatment, cells were treated with 500 μg/mL spectinomycin to stop protein synthesis and incubated at 30 °C. Immediately after spectinomycin treatment and 2 hours after stopping protein synthesis, respectively, aliquots of cells were placed on 1.5% agarose phosphate buffer pads to visualize aggregate formation and clearance. The quantification of aggregate numbers in each cell was done manually in ImageJ.

This protocol was altered for tracking aggregate clearance in single cells. Overnight cultures were additionally grown in ampicillin if the cells contained the pZS*11-P_Tet_-*mCherry* plasmid. Once the cultures reached mid-logarithmic phase, they were incubated at 42 °C for 30 minutes. After treatment, cells were treated with 500 μg/mL spectinomycin to stop protein synthesis and immediately transferred to a 200 μl agarose LB pad containing 500 μg/mL spectinomycin. Then fluorescence and DIC images were taken immediately and at 3 hours. The quantification of aggregate numbers was done manually in ImageJ, XXX cells in one image and three different images were selected for qualification.

### Determination of RpoH protein level

Overnight cultures were diluted 1:100 or 1:50 (Δ*rpoS* strains) in LB and grown for 3 hours in LB at 30 °C. 1 mL of the culture was treated with 100 μL of cold trichloroacetic acid (TCA), and then incubated on ice for at least 10 minutes. Samples were centrifuged at 13,000 rpm for 10 minutes at 4 °C. After discarding the supernatant, the pellet was washed in 500 μL of 80% cold acetone. The samples were centrifuged and the supernatant discarded. The samples were dried in open air at room temperature for 30 minutes, then resuspended in 50 μL of 7 M urea to resuspend the pellet. To facilitate complete resuspension of the pellet, the samples were incubated at 95 °C while shaking. Once the pellet was completely resuspended, the samples were stored at -80 °C.

The lysate protein concentration was determined via bicinchoninic acid assay (BCA assay) according to manufacturer’s directions (ThermoFisher Scientific. The RpoH protein level was determined using Western blotting.

### Gene Expression analysis via *E. coli* Promoter Collection strains

Strains containing a low-copy number plasmid with selected promoters fused to *gfp* were grown at 30°C shaking in a 96-well plate and fluorescent measurements taken every 20 minutes. The values used for analysis were at the timepoint where the OD600 was closest to 0.4. For data analysis, expression of the RpoH-target promoters was normalized to *ppa* expression to account for potential changes to overall transcription levels between MG1655 and *rpsD\**.

## Results

### Increased mistranslation protects against heat stress

Increased levels of mistranslation have been shown to have a protective effect on cells by pre-activating stress responses that are beneficial when a population encounters lethal stress [23, 24]. Mistranslation results in the production of misfolded proteins; therefore, we hypothesized that increased levels of mistranslation would protect populations from heat stress by pre-activating the heat shock response. To model cells experiencing a high mistranslation rate, we used the error-prone *E. coli* strain, *rpsD\**. This strain contains a ribosomal point mutation, I199N, which decreases ribosomal accuracy [17, 25].

To determine whether mistranslation protects against heat stress, we performed a heat killing assay with *rpsD\** and its parental wild type strain, MG1655. In this assay, cells were grown to mid-logarithmic phase in LB then subjected to killing at 50 °C. Cell survival was assayed by counting colony forming units. The *rpsD\** strain survived heat killing better than MG1655 (Figure 1). This led us to hypothesize that *rpsD\** pre-activates the heat shock response during normal growth. To test this, we measured the activation of transcriptional reporters with RpoH-dependent promoters fused to GFP. We found that the expression of two major RpoH targets, *dnaK* and *groE*, is 2-fold higher in *rpsD\** than in MG1655; however, the transcription of *rpoH* was unaffected in the *rpsD\** strain (Figure 1).

**Figure 1:**
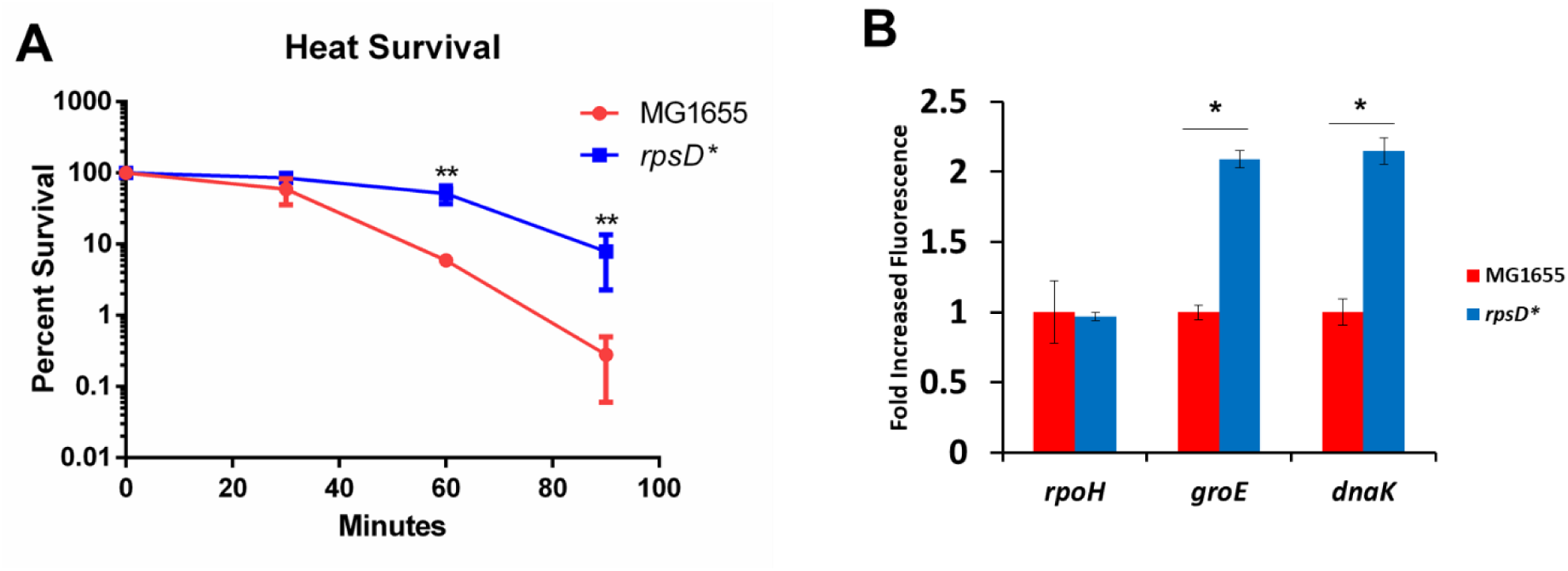
The *rpsD*\* strain is protected from heat killing and has an activated heat shock response. (A) MG1655 and *rpsD*\* strains were grown to mid-logarithmic phase at 30 °C, then treated at 50 °C for 90 minutes. Serial dilutions were made every 30 minutes and CFU counts were used to determine the rate of killing (N=3). (B). MG1655 and *rpsD*\* strains expressing promoter fusions to *gfp* on a low copy plasmid were grown to mid-logarithmic phase. The fluorescence in each culture was determined using fluorescence spectroscopy (N=3). *p-value < 0.05. **p-value < 0.01 by unpaired t-test.

In addition to heat killing, we determined whether *rpsD\** cells have an advantage in non-lethal protein misfolding stress conditions. To do this, we created a fluorescent reporter, sfGFP-ClpB. This reporter is a fusion between superfolder-GFP (sfGFP) and the primary protein chaperone disaggregase, ClpB [26]. With this reporter, protein aggregates formed in cells can be visualized as green fluorescent foci as ClpB binds to the aggregates. We used this reporter to determine the ability of cells to clear aggregates after being exposed to protein misfolding stresses – non-lethal heat or streptomycin treatment-by visualizing the clearance of aggregates after stress treatment.

In response to incubation at the non-lethal heat stress, 42 °C, both MG1655 and *rpsD\** formed aggregates in every cell (Figure 2). The number of aggregates formed were visually quantified and the *rpsD\** strain formed fewer aggregates per cell. In comparison, after recovering from heat stress for 2 hours there was a dramatic difference in the number of aggregates in *rpsD\** cells and MG1655 cells. On average, MG1655 cells still had ∼2.5 aggregates per cell, whereas *rpsD\** cells had cleared most aggregates and had only ∼0.5 aggregates per cell after 2 hours of recovering (Figure 2 A and B). This assay was repeated with the antibiotic stress, streptomycin treatment. This aminoglycoside antibiotic binds directly to the ribosome and drastically increases mistranslation, resulting in the accumulation of misfolded proteins. After recovery from streptomycin treatment, *rpsD\** cells were able to clear streptomycin-induced aggregates more effectively than MG1655 cells (Figure 2 C and 2D).

**Figure 2:**
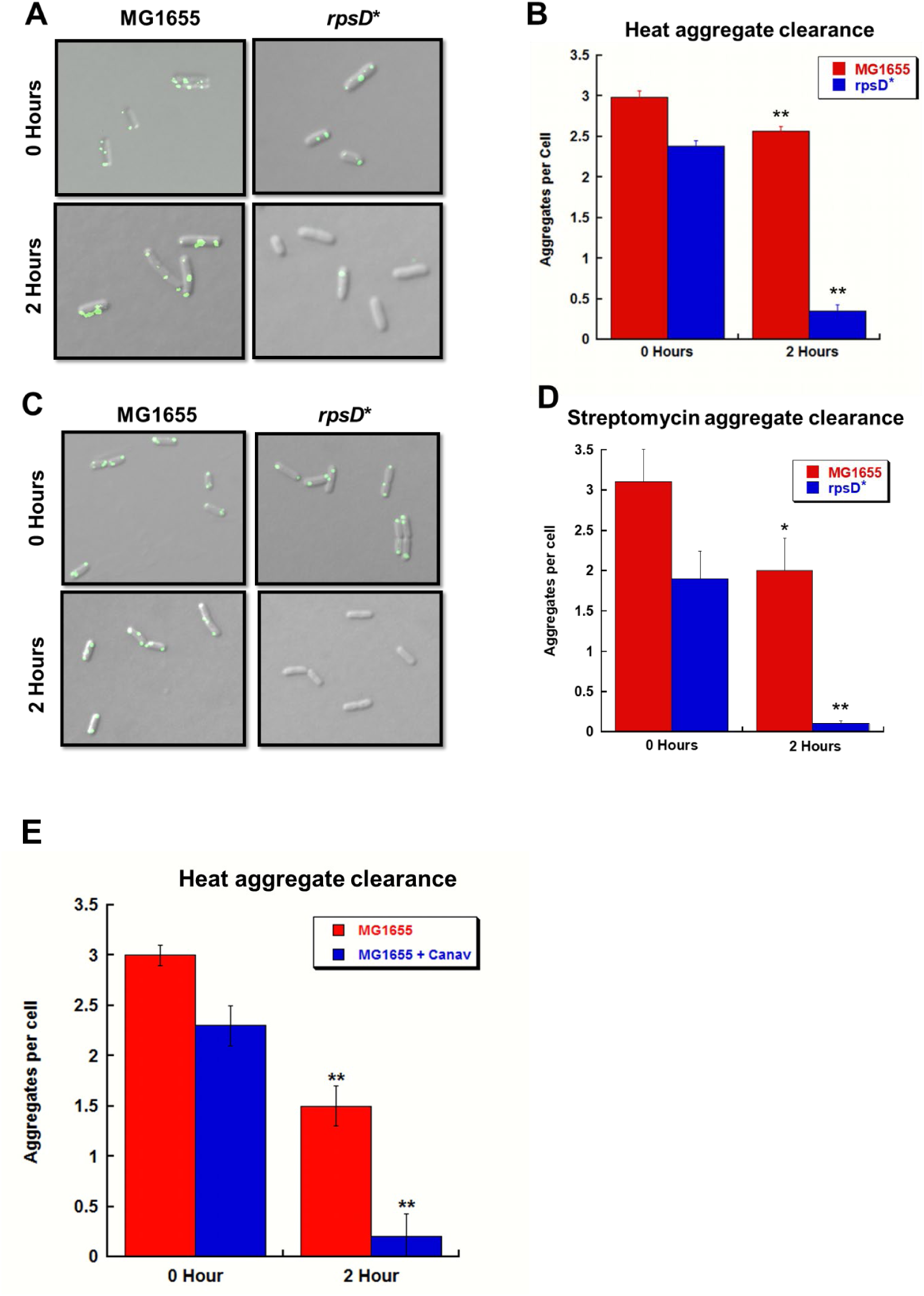
Increased mistranslation induces aggregate clearance. MG1655 and *rpsD*\* strains expressing an IPTG-inducible sfGFP-ClpB fusion were grown to mid-logarithmic phase in LB with 100 μM IPTG at 30 °C. Then, cells were treated at 42 °C (A and B) or with 100 μg/mL streptomycin (C and D) for one hour. After treatment, cultures were treated with 100 μg/mL spectinomycin to stop protein synthesis. Images were taken immediately after and two hours after spectinomycin treatment. (A and C) Representative images of cells after treatment. (B and D) The numbers of aggregates per cell were quantified manually using imageJ (N=3). (E) MG1655 cells expressing sfGFP-ClpB were grown in the presence or absence of the arginine analogue, Canavanine (Canav), to mid-logarithmic phase in LB with 100 μM IPTG. After growth, cultures were treated at 42 °C for one hour. After treatment, cultures were treated with 100 μg/mL spectinomycin to stop protein synthesis. Images were taken immediately after and two hours after spectinomycin treatment. The numbers of aggregates per cell were quantified manually using ImageJ (N=3). *p-value < 0.05, **p-value < 0.01 by unpaired t-test compared to time point 0 hours.

To ensure that the increased disaggregation is not a *rpsD\** strain-specific phenotype, we tested whether increasing mistranslation in MG1655 via canavanine treatment during growth would recapitulate the protective phenotypes seen in *rpsD\**. Canavanine is an arginine analogue that is misincorporated by the ribosome into proteins in place of arginine and results in the production of proteins that misfold [27]. MG1655 cells expressing the sfGFP-ClpB reporter were grown with or without canavanine treatment and treated at 42 °C, then their ability to clear aggregates was quantified. As with the *rpsD\** strain, MG1655 cells grown in canavanine were able to clear heat-induced aggregates from cells better than MG1655 cells grown without canavanine (Figure 2E).

In the experiments described above, samples of the population were assayed at different time points to determine how the population was recovering from stress. We aimed to visualize single cells recovering from stress immediately after treatment until they had fully recovered. This allows us to more accurately determine how cells were able to recover and the differences between MG1655 and *rpsD*\* cells. To accomplish this, we expressed mCherry from a constitutive tetracycline-on promoter in either MG1655 or *rpsD\** strains along with the sfGFP-ClpB reporter. Then, after growth and heat stress, we visualized the recovery of MG1655 and *rpsD*\* cells together on an LB agarose pad and directly compare their recovery rates (Figure 3A). Using this method, we found that over 90% of *rpsD*\* cells were able to completely clear their heat-induced aggregates in 3 hours, while ∼70% of MG1655 cells still had at least one aggregate in the same timeframe (Figure 3B).

**Figure 3:**
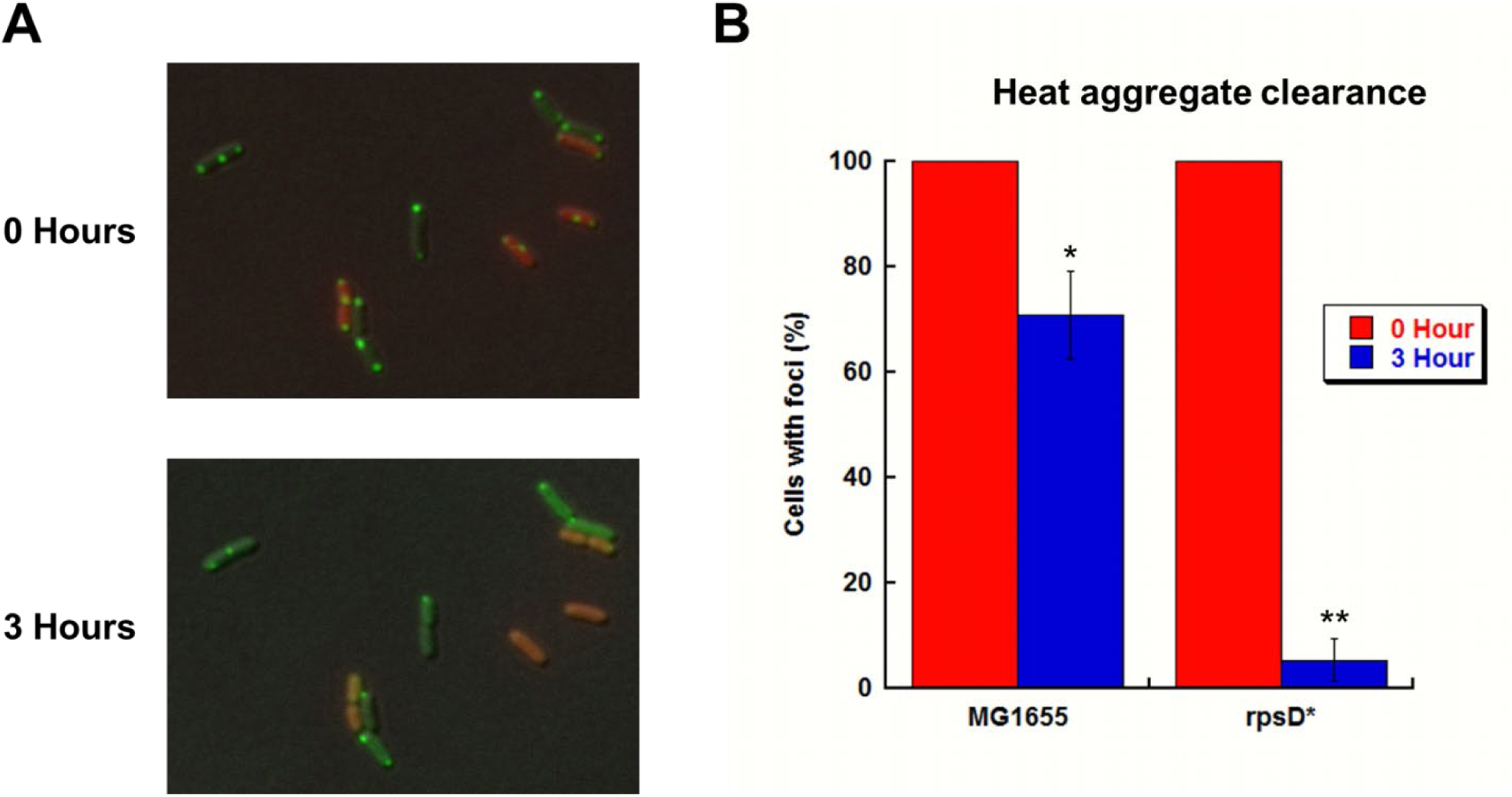
Time-lapse microscopy reveals that increased mistranslation promotes aggregate clearance in single cells. MG1655 and *rpsD*\* cells expressing the IPTG-inducible sfGFP-ClpB construct were grown in LB with 100 μM IPTG to mid-logarithmic phase. Additionally, the *rpsD*\* cells constitutively expressed mCherry on a plasmid. After growth, cultures were mixed and incubated at 42 °C for 30 minutes. Then, spectinomycin were added to the culture to stop protein synthesis. Cells were transferred to a 1.5% agarose pad. (A) Aggregate formation was visualized using fluorescence microscopy and recover was tracked for 3 hours. (B) The number of cells containing aggregates in the MG1655 and *rpsD*\* cells were quantified manually using ImageJ (N=2 of >60 individual cells). *p-value < 0.05, **p-value < 0.01 by unpaired t-test.

### The heat shock response is pre-activated in *rpsD\**

We hypothesized that the benefits to the *rpsD\** cells against misfolded protein stress was due to the pre-activation of the heat shock response. We measured the levels of the heat shock sigma factor, RpoH, under normal growth conditions to determine whether mistranslation was sufficient to activate the heat shock response in *rpsD\**. To measure RpoH levels, we performed Western blotting on lysates using an anti-RpoH antibody. Under normal growth conditions, *rpsD\** increases RpoH protein levels over 2-fold higher than in MG1655 cells (Figure 4A, lanes 1 and 2). This appears to be an incomplete activation of the heat shock response, as MG1655 cells under heat conditions increase their RpoH levels over 5-fold (Figure 4A, lane 4).

**Figure 4:**
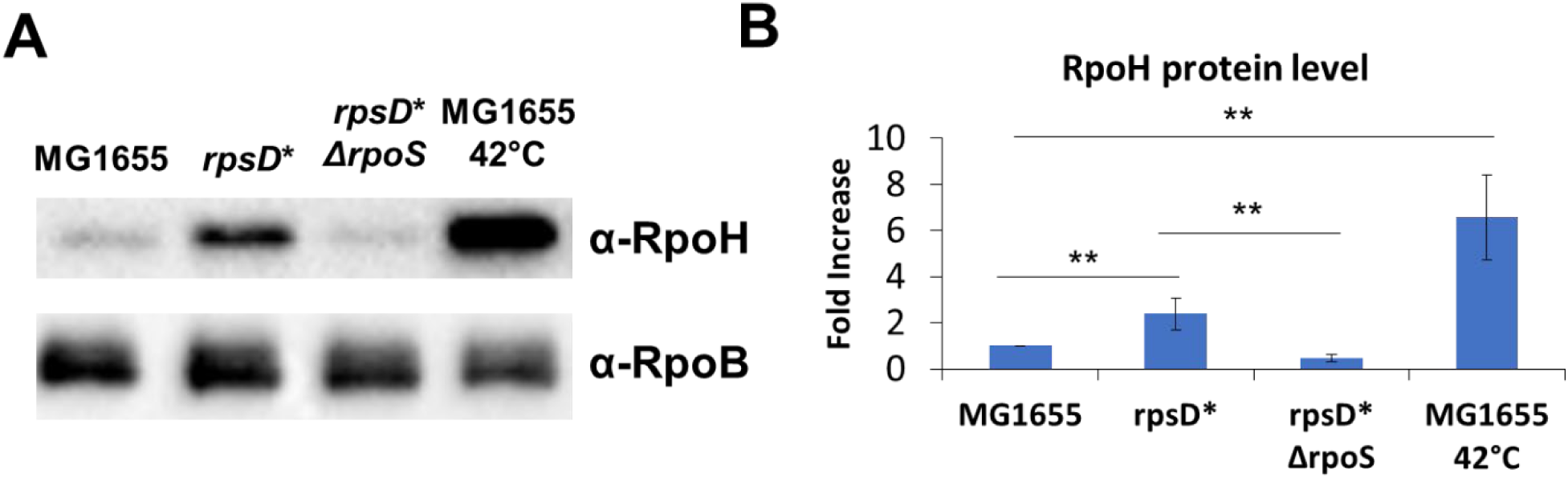
RpoS is necessary for the mistranslation-induced increase of RpoH. MG1655, *rpsD*\*, *rpsD*\* Δ*rpos* strains were grown to mid-logarithmic phase in LB at 30 °C. For comparison to RpoH induction by heat shock, MG1655 cells were incubated at 42 °C for 20 minutes. (A) A western blot of cell lysates using an α-RpoH antibody to determine RpoH levels in each strain. RpoB protein levels were determined as a loading control. (B) Three independent repeats were quantified using volumetric analysis. **p-value < 0.01 by unpaired t-test.

### RpoS is necessary for the protective mistranslation-induced heat shock response

The levels of the general stress response sigma factor, RpoS, is increased in the *rpsD*\* strain [17]. The increase in RpoS is responsible for the protective effect of mistranslation against oxidative stress [17]. Additionally, there is some evidence that RpoS could play a role in the protection of cells from heat stress, although no conclusive mechanism has been identified [28].

We tested whether RpoS plays a role in the survival of *rpsD*\* in 50 °C lethal heat treatment. The deletion of *rpoS* in the *rpsD\** background (*rpsD\** Δ*rpoS*) decreases the heat survival of *rpsD\** (Figure 5A). This phenotype is not *rpsD*\*-specific. MG1655 grown with canavanine survives lethal heat stress better than MG1655 grown without canavanine (Figure 5B). Consistent with our model, canavanine does not protect MG1655 Δ*rpoS* cells the same as MG1655 against heat killing (Figure 5B).

**Figure 5:**
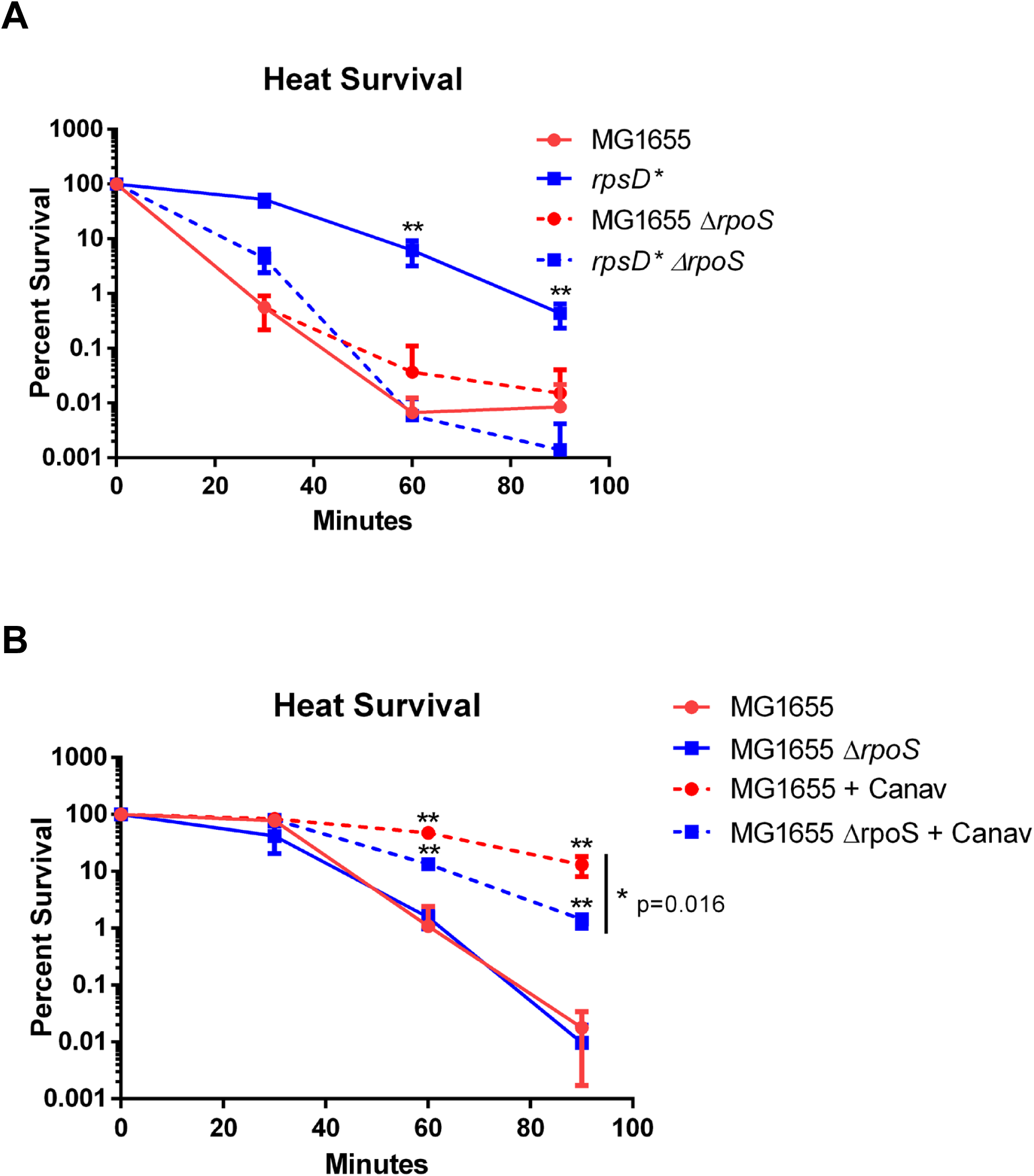
Mistranslation-induced heat protection is dependent on RpoS. (A) *rpoS* deletion strains of MG1655 and *rpsD*\* were grown to mid-logarithmic phase at 30°C, then incubated at 50°C. Heat killing was assayed via colony formation by serial dilution every 30 minutes. (B) The same experiment was performed with MG1655 with or without *rpoS* grown in the presence of canavanine (Canav). N≥3 **p-value < 0.01 by unpaired t-test for each point compared to MG1655.

To determine whether RpoS affects the activation of the mistranslation-induced heat shock response, we determined whether the presence of RpoS affects the production of RpoH in the *rpsD*\* strain. In cells grown to mid-logarithmic phase, the RpoH protein level of *rpsD\** is ∼2 fold higher than in MG1655 cells. In the *rpsD\** Δ*rpoS* strain, the RpoH level decreases to the level of MG1655 (Figure 4A). This indicates that mistranslation-induced RpoS increases heat protection by affecting heat shock response activation.

## Discussion

We used two model systems to examine the physiological effect of mistranslation, an endogenous source of mistranslation via ribosomal mutation and an exogenous source via treatment with an antibiotic, canavanine. Using these model systems, we found that increasing the mistranslation rate above wild-type levels resulted in protection from heat killing. This led us to a model by which cells may alter their mistranslation rate in order to increase their survival in conditions which may otherwise be lethal. This benefit of increased translation error rate may help explain the observation that ribosomal mutations that increase mistranslation are common in natural isolates of *E. coli* [13]. It is interesting to consider that the ribosome and protein synthesis machinery may act as internal stressors that can prepare cells for future stresses. The heat protection is another benefit to add to the growing list of mistranslation-induced benefits including oxidative stress resistance [17], antibiotic resistance [15], and facilitating more efficient adaptive evolution [29].

During heat shock, the production of RpoH is independent of RpoS; however, the activation of RpoH in the absence of heat is not well understood. In response to misfolded protein production, the presence of oxygen is required for RpoH activation, indicating that solely the production of misfolded proteins, which should result in an increase in RpoH stability via sequestration of the protease machinery, is not sufficient to activate the response [30]. Here, we show that RpoS is necessary to increase RpoH protein levels in response in increased mistranslation rates. It is appealing to hypothesize that oxidative damage to misfolded proteins results in the activation of RpoS, which is well-known to be involved in the oxidative stress response. The activation of RpoS in combination with the accumulation of misfolded proteins and sequestration of chaperones and proteases is then sufficient to activate the RpoH heat shock response. Further, the mechanism by which RpoS increases the levels of RpoH remains a mystery. Direct transcriptional regulation by RpoS is unlikely since we were unable to detect any transcriptional changes to *rpoH* in the presence or absence of RpoS.

In wild isolates of *E. coli*, purified ribosomes have been found to have a range of accuracy [13]. However, upon growth and selection within a laboratory setting, the strains mutated their ribosomes to have an accuracy rate comparable to a known laboratory strain. Clearly, evolutionary pressure can impact ribosome activity and accuracy to broadly affect cell physiology to adapt to changing environmental conditions. Our study supports a model that ribosomal mutations that decrease accuracy in natural isolates may increase strain fitness by protecting cells from environmental stresses, such as increased heat or conditions that stress the proteome by increasing misfolding.

## Acknowledgements

The authors would like to thank the members of the DeLay and Kim labs for their technical assistance and input. This work was funded by National Institute of General Medical Sciences [grant number R01GM115431] to J.L.

